# Evidence for Bias of Genetic Ancestry in Resting State Functional MRI

**DOI:** 10.1101/440776

**Authors:** Andre Altmann, Janaina Mourao-Miranda

**Affiliations:** Centre for Medical Image Computing, University College London, UK; Max Planck University College London Centre for Computational Psychiatry and Ageing Research University College London, UK

**Keywords:** resting state fMRI, genetics, ancestry, imaging genetics, machine learning

## Abstract

Resting state functional magnetic resonance imaging (rs-fMRI) is a popular imaging modality for mapping the functional connectivity of the brain. Rs-fMRI is, just like other neuroimaging modalities, subject to a series of technical and subject level biases that change the inferred connectivity pattern. In this work we predicted genetic ancestry from rs-fMRI connectivity data at very high performance (area under the ROC curve of 0.93). Thereby, we demonstrated that genetic ancestry is encoded in the functional connectivity pattern of the brain at rest. Consequently, genetic ancestry constitutes a bias that should be accounted for in the analysis of rs-fMRI data.

## 1. INTRODUCTION

Task free or resting state (rs) functional magnetic resonance imaging (fMRI) is an increasingly popular modality to map the functional connectivity pattern of the brain. Recent rs-fMRI applications include functional brain parcellation [1] and disease biomarkers [2]. However, rs-fMRI is subject to various biases that alter the inferred functional connectivity and may obscure true disease effects or genuine brain function. For instance, observed connectivity patterns can vary strongly with scanner model, pulse sequence and scan site [3]. Also, involuntary head movement of the subject during the scan can induce spurious functional connectivity between brain regions [4]. In addition to these technical biases, there are also biases originating from subject level characteristics: subjects’ sex and age have been reported to exhibit wide-spread effects on the observed resting state connectivity pattern [3]. In fact, subjects’ age can be successfully predicted from rs-fMRI [5]. Furthermore, the involuntary changes in cognitive states and vigilance levels has profound changes in functional brain connectivity [6].

These biases are often taken into consideration when planning rs-fMRI studies. Typically, technical biases are minimized by limiting studies to a single MRI scanner and by addressing known artifacts, e.g., the ones arising from head movement. Subject level characteristics are either considered by matching the distributions, e.g., of age and sex, between the studied disease groups or by including these subject characteristics as confounding variables in the statistical analysis. However, there is one additional subject level characteristic that is known to affect head and brain morphology but is rarely considered as a confound in rs-fMRI studies or brain imaging studies in general: *genetic ancestry*.

The human genome was in part shaped by mankind’s migration history across the globe. Statistical analysis of genetic data can reveal these ancient migration patterns. For instance, the two main principal axes of variation in genetic similarity of European individuals reflect the north-south and east-west gradient within Europe [7]. The same principle holds true for differences between continental regions across the world. Thus, it is possible to reliable extract ancestry information from genetic data, which is referred to as genetic ancestry and mostly reflects subjects’ self-reported ethnicity.

Previous work on the relation of genetic ancestry and brain imaging has demonstrated that head and brain morphology of people with European ancestry follow the same north-south and east-west pattern [8]. Follow-up work demonstrated that the human cortical surface encodes the genetic ancestry and that regional patterns of cortical folding and gyrification are unique and complex for each continental ancestry [9].

In this paper we demonstrate that functional connectivity networks obtained from high-quality rs-fMRI can reliably predict genetic ancestry derived from genome wide genotyping data. In section 2 we describe the data used for this work as well as the statistical analysis. Section 3 summarizes the results which are discussed in section 4.

## 2. MATERIAL AND METHODS

### 2.1. Genetic and imaging data

We obtained rs-fMRI data and matched genetic data from the Young Adult study of the Human Connectome Project (HCP) [10]. The HCP aims at charting the neural pathways that underlie brain function and behavior and has acquired high-quality neuroimaging data in over 1,100 healthy young adults aged 22 - 35. Behavioral and other individual subject measure data are available on all subjects.

#### 2.1.1. Imaging data

The HCP acquired four imaging modalities using a 3T MR for all subjects: structural images (T1w and T2w), resting-state fMRI (rs-fMRI), task-fMRI (tfMRI), and high angular resolution diffusion imaging (dMRI). The rs-fMRI data were acquired in four runs of approximately 15 minutes each, two runs in one session and two in another session, with eyes open with relaxed fixation on a projected bright cross-hair on a dark background (and presented in a darkened room). The TR was 0. 75s amounting to 4,800 time-points per subject.

Detailed description of acquisition parameters and processing steps can be found in [11]. In brief, group-average parcellations for all subjects were obtained using group-ICA at several different dimensionalities (15, 25, 50, 100, 200, 300). Next, subject-specific sets of node time series were extracted, where each ICA-defined region of interest (ROI) acted as a node. Then for each subject, a node × node connectivity matrix was created by computing the temporal correlation between each pair of nodes. Correlation estimates were based either on Pearson’s correlation or partial correlation using Tikhonov regularization. For the analyses presented here we obtained the connectivity matrices based on partial correlation at each of the six resolution levels.

The subjects’ matrices were vectorized to render them amendable for machine learning; we extracted the upper right triangle (omitting the diagonal values) of the correlation estimates, thus generating for each subject feature vectors, *x*, of length 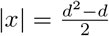, where *d* is the number of input ROIs.

#### 2.1.2. Genetic data

Genetic data in HCP were available for 1,141 participants. For this work we accessed the genome-wide genotyping data that measured the identity of 2,119,803 genetic variants. In brief, genotyping arrays assess the identity of genetic variants at predefined locations in the genome. A variant can either be identical to the one listed in the human reference genome (*reference*) or differ from the reference (*alternative*). Humans carry two copies of the genome in each cell. For analysis purposes, one simply counts for each tested genetic variant the number of alternative copies. Thus, at each of the two million positions either 0, 1 or 2 is recorded.

### 2.2. Estimation of genetic ancestry

We used SNPweights [12] to obtain predictions for genetic ancestry for four continental groups: Europeans (`CEU`), African (`YRI`), Asian (`ASI`) and Native American (`NAT`). SNPweights provides for each subject, *i*, and each continental group, *g*, a probability 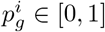. Furthermore, probabilities for each subject sum to 1 for all *N* subjects:

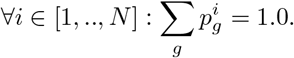

### 2.3. Statistical learning

Our objective was to predict genetic ancestry from rs-fMRI connectivity data. We employed the elastic net classifier [13], which uses both the 𝓁_1_ and 𝓁_2_ penalties during training:

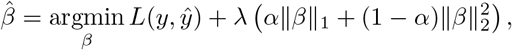

where *y* is the vector of target values, *β* is a vector of model coefficients for each feature, 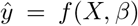 is the vector of model predictions, *X* is the feature matrix, 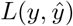 is the loss function, ||*β*||_1_ = Σ|*β_j_*| is the 𝓁_1_ penalty, and 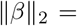 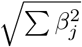 is the 𝓁_2_ penalty. The loss function is typically either the squared loss for regression problems or the logistic loss for classification problems. The *α* parameter trades off the 𝓁_1_ and 𝓁_2_ penalties. Of note, *α* = 0 corresponds to ridge regression and *α* = 1 corresponds to the least absolute shrinkage and selection operator (LASSO) regression. Thus, for settings of *α* other than 0, the elastic net produces a sparse solution with many entries in 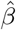 set to 0. In practice, the exact choice of *α* (other than at the two extremes) has limited impact on model performance. Therefore, we set *a priory α* = 0.5. A more crucial parameter is λ, which trades off the the amount of regularization with the model fit to the training data. Larger values of λ tend to result in models with more entries in *β* being set to 0 (i.e., sparser models). The λ parameter is typically optimized using cross-validation (CV). For the experiments described in this work we used the elastic net implementation in the R package `glmnet`.

Depending on the selected resolution, each subjects’ feature vector ranged from 105 to 44,850 entries for 15 and 300 ROIs, respectively.

The target vector *y* was based on the genetic ancestry prediction. We dichotomized the quantitative output in order to be able to perform classification rather than regression. One consideration was that the dataset is rather imbalanced (see section 3) and the target values ranged from 0.0 to 1.0 and are therefore likely to suffer from the floor and ceiling effects. To this end we employed two cutoffs for *p*_CEU_: 0.5 constituting “mainly European” and 0.9 constituting “predominantly European”.

Classifier performance was assessed using receiver operating characteristics (ROC) curves and the area under the ROC curve (AUC). We used nested CV with 10 outer folds and 5 inner folds. In nested CV, the data is first split into outer folds (10 here). Then, data from all-but-one outer fold are used to train the classifier. The classifier’s parameters (λ here) are optimized through the inner CV (5-fold CV here). The resulting classifier is then used to predict the label for the data in the left-out outer fold to obtain an unbiased performance estimate. The process is repeated until each outer fold served once as hold-out data. Within the inner CV, we selected the largest value of λ (i.e., the sparsest model) such that the AUC was within 1 standard error of the maximum AUC achieved by all tested λs; this strategy (“1 standard-error rule”) is commonly employed to minimize the risk of overfitting by selecting a sparser model that does not perform significantly worse than the best model [13]. Of note, the HCP data contains data from siblings, which have a high genetic similarity. In order to avoid confounding the performance estimate in cases where one sibling contributes to training and the other sibling contributes to the testing data, we have sampled the outer CV folds such that all family members were placed within the same fold. In addition, the sampling for the the outer fold was retained for all experiments to ensure that results are comparable between settings.

In order to put the AUC results into the broader context we also predicted subjects’ sex from the same rs-fMRI data with the same CV folds using the 300 ROI resolution.

## 3. RESULTS

There were 1003 subjects with processed rs-fMRI data available through HCP, of these, 950 subjects (502 females) also had genome-wide genotyping data available. SNPweights successfully produced genetic ancestry predictions for the four continental groups for all 950 subjects. When considering the highest probability for predicted genetic ancestry (i.e., “argmax”), then HCP comprises 764 `CEU`, 138 `YRI`, 39 `ASI` and 9 `NAT` subjects. Figure 1 depicts the distribution of genetic ancestry predictions restricted to the three most predominant groups in HCP: `CEU`, `YRI` and `ASI`. It becomes evident that not only the corners of the triangle are populated, i. e., representing uniform genetic ancestry, but there are also many subjects with mixed ancestries, referred to as *genetic admixture*.

As rationalized in section 2, we dichotomized the genetic ancestry information for classification purposes. Given that the majority of the HCP participants were of European ancestry, we set a cutoff at *p*_CEU_ > 0.5 resulting in 748 `CEU` and 202 non-`CEU` subjects and the other second more restrictive cutoff at *p*_CEU_ > 0.9 resulting in 651 `CEU` and 299 non-`CEU` participants. We then employed the elastic net classifier to predict `CEU` status using the vectorized rs-fMRI connectivity matrices. Practically, we were aiming to classify subjects who are in the lower right corner of the triangle in Figure 1 versus subjects who are not, based on their rs-fMRI connectivity. Table 1 summarizes the mean and standard deviation of AUC values for the 10 outer folds of the CV for both genetic ancestry cutoffs and all six levels of resolution. Figure 2 depicts representative ROC curves for selected ROI resolutions at the *p*_CEU_ > 0.5 cutoff. AUC values ranged from 0.72 to 0.93; models using a finer-grained cortical parcellation showed higher performance despite a high features to samples ratio of ≈ 47 when using connectivity matrices based on 300 ROIs.

**Fig. 1.**
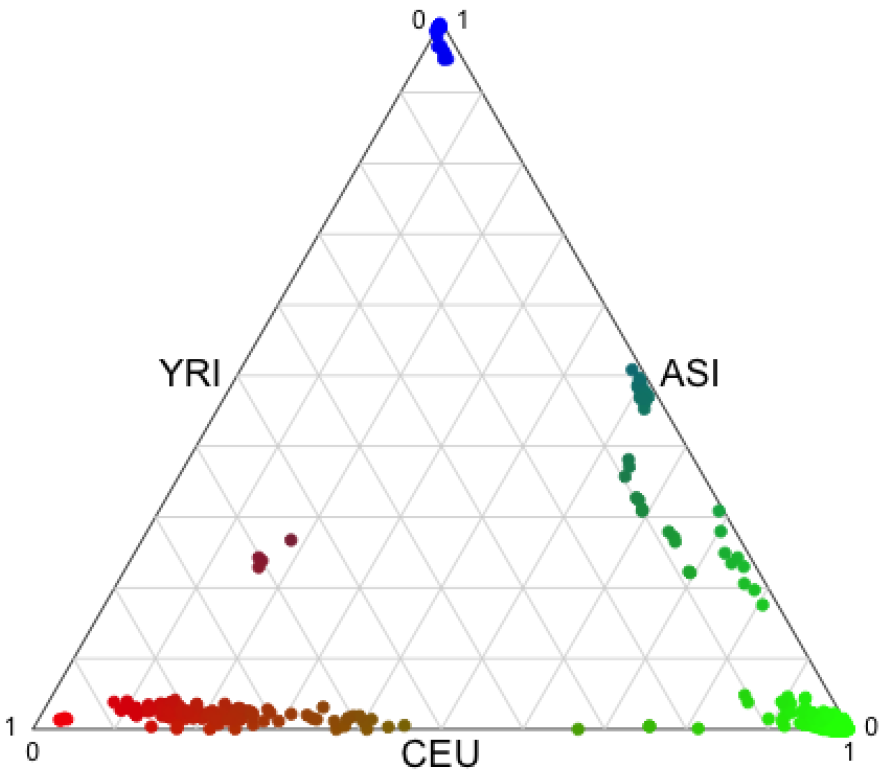
Distribution of the three main genetic ancestries in the HCP dataset. 80 subjects with *p*_NAT_ > 0.05 had been omitted from this plot.

Subjects’ sex could also be predicted at very high accuracy with mean AUC of 0.98 (and standard deviation of 0.016).

**Table 1.** Classifier performance quantified by mean AUC and standard deviation from 10 × 5-fold nested CV for different settings of *p*_CEU_ cutoff and number of ROIs.

| ROIs | dim | $p_{\text{CEU}} > 0.5$ | $p_{\text{CEU}} > 0.9$ |
| --- | --- | --- | --- |
| 15 | 105 | 0.78 (0.088) | 0.72 (0.049) |
| 25 | 300 | 0.81 (0.060) | 0.76 (0.061) |
| 50 | 1225 | 0.86 (0.055) | 0.83 (0.063) |
| 100 | 4950 | 0.91 (0.046) | 0.86 (0.061) |
| 200 | 19900 | 0.92 (0.039) | 0.88 (0.055) |
| 300 | 44850 | 0.93 (0.036) | 0.87 (0.032) |

**Fig. 2.**
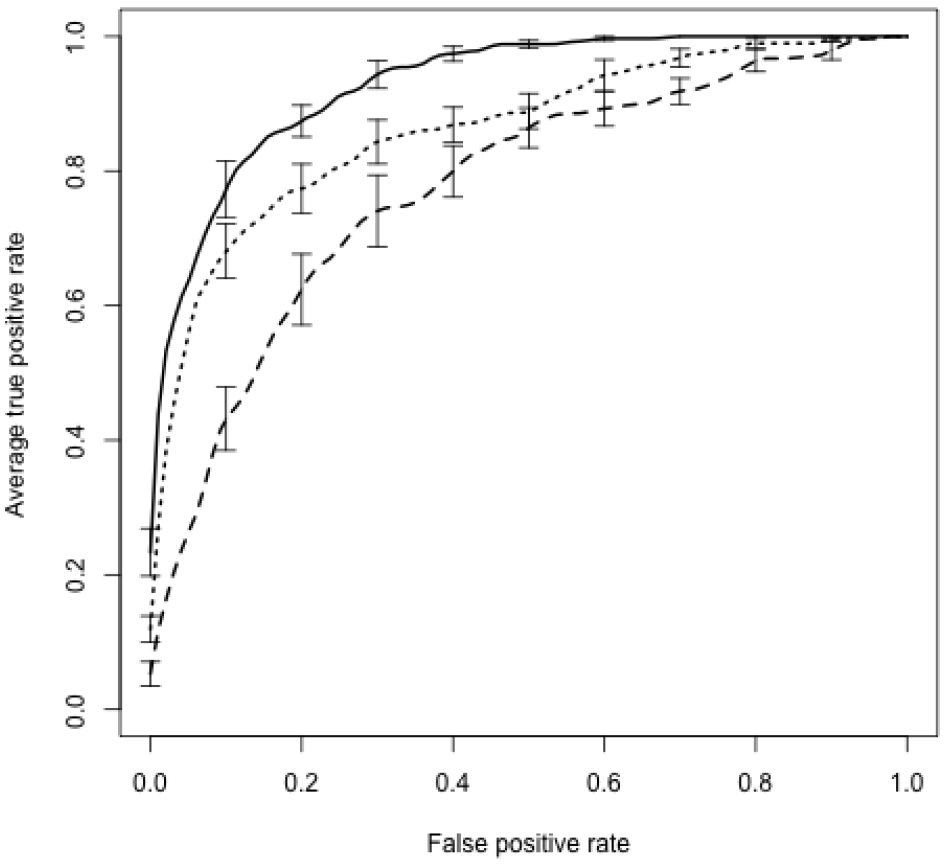
Averaged ROC curves from the 10 outer CV-folds. Class labels were based on the *p*_CEU_ > 0.5 cutoff. Dashed, dotted and solid lines correspond to models based on 15, 50 and 300 ROIs, respectively. Whiskers indicate 1 standard error estimates.

## 4. DISCUSSION

This is the first work demonstrating that genetic ancestry is highly predictable from rs-fMRI connectivity patterns. Our results indicate that genetic ancestry is a serious bias that modifies estimated brain connectivity and may mask genuine differences or may introduce spurious differences in rs-fMRI analyses between groups, e.g., a disease group and a control group. The extent of the bias is not as pronounced as the influence of participants’ sex, which showed near perfect classification from rs-fMRI.

The exact origin of these apparent connectivity differences between continental ancestries remains elusive at the moment. However, we hypothesize that the observed differences are not based on true neuronal differences but that they originate from differences in head and brain morphology as reported in [8, 9]. These morphological differences may be carried forward through the standard rs-fMRI processing pipeline and affect the inferred functional connectivity. In addition, rs-fMRI connectivity is based on correlations between blood-oxygen-level dependent (BOLD) signal time series at rest. Thus, it is conceivable that genetic differences contributing to blood circulation, perfusion and elasticity of the vascular system may modify BOLD dynamics. This is exemplified by reports identifying ethnicity as independent risk factors for cardiovascular disease [14] and intracranial artery tortuosity [15]. In addition, brain hemodynamic responses are known to be heritable traits [16].

This study is not without limitations. Firstly, we chose to dichotomize genetic ancestry and to perform classification rather to train a regression model; this was mainly owing to sample imbalance and to anticipated floor and ceiling effects when working with probability scores. Secondly, in this proof-of-principle study we only classified European (`CEU`) individuals from non-European individuals; studies on larger datasets would be able to build classifiers for each continental ancestry. Lastly, the analysis was limited to continental ancestries; with sufficiently large datasets it should be possible to assess the effect of sub-ancestries within one continental group on rs-fMRI inferred connectivity.

This work exemplifies that in the domain of rs-fMRI analysis there is a need to consider genetic ancestry as a confound in the analysis. Given the previous findings of the strong influence of genetic ancestry on regional volume and gyri-fication [9], this consideration may extend to the entire neuroimaging field.

